# Deriving microbial food web structure by maximizing entropy production over variable timescales

**DOI:** 10.1101/2025.04.30.651452

**Authors:** Joseph J. Vallino, Olivia M. Ahern, Julie A. Huber

**Affiliations:** Ecosystems Center, Marine Biological Laboratory, Woods Hole, MA; Marine Chemistry and Geochemistry, Woods Hole Oceanographic Institution, Woods Hole, MA

**Keywords:** Maximum entropy production, Food web, biogeochemistry, trait-based modeling

## Abstract

Protists and viruses dynamically alter the flow of mass and energy through microbial food webs via predation. Simple microbial food web models show that the addition of microbial predators can increase the primary production of a microbial community but only for some configurations of food web structure. Under the conjecture that systems self-organize to maximize energy dissipation, known as the maximum entropy production (MEP) principle, we developed an MEP-based model that predicts microbial food web structure, and we examine how food web structure differs when entropy production is maximized over short versus long timescales. The model design follows from an experimental system and uses a trait-based variational method to set trait values by maximizing entropy production over a specified interval of time. Model results show that short-term MEP optimization (STO) produces microbial communities that specialize in substrate preference and consumers that have fewer trophic levels than solutions based on long-term optimization (LTO) that have substrate generalist and more trophic levels. Our MEP-based approach provides an alternative to food web structure synthesis that does not depend on assumptions of community stability.

## Introduction

The Workshop on Information, Selection, and Evolution (WISE), which is the theme of this *Interface Focus* issue, entangled scientists and philosophers in discussions on evolution, emergence, selection, function, information, energy/entropy, statistical mechanics, and the arrow(s) of time. The maximum entropy production (MEP) principle can be used to predict the structure and function of microbial food webs, such as marine planktonic ecosystems that supply energy and nutrients to nearly all ocean life [1], as well as the Earth’s microbiome that governs the cycling of elements from local to global scales (aka, biogeochemistry) [2]. The MEP principle follows from statistical mechanics applied to non-equilibrium steady-state systems and proposes that systems with many degrees of freedom will *likely* organize to maximize the rate of entropy production, or equivalently, the rate of free energy dissipation [3-8]. The theory is agnostic on the type of system and, while it has been applied to many abiotic processes [9, 10], its application to biological systems may provide a directionality to evolution [11-16]. In theoretical ecology, there is a long history of proposed principles that attempt to describe how living systems evolve, organize and function, but many of the ideas only apply to biological systems or are based on ad hoc principles. Some of the early theories are thermodynamically inspired, and these have recently been formalized under the MEP principle [17, 18], provided there is some relaxation to the steady-state constraint that the formal theory currently is limited to. When time is explicitly included in the problem, a distinction can be made between living (biotic) and non-living (abiotic) systems. In particular, abiotic systems maximize entropy production instantaneously, while biotic systems maximize entropy production integrated over time [15, 19]. While this is an important concept that will be briefly revisited in the Methods Section, the theory has been developed in other studies [15, 20]. Here, we extend the MEP approach to predict how the connectivity of microbial foods webs (i.e., who eats whom) can be predicted by maximizing entropy production over time.

The structure of food webs, at both the microscale and macroscale, dramatically impacts community dynamics as well as how energy and nutrients are used and processed at the system level [21]. For instance, even though herbivores eat plants (and zooplankton eat phytoplankton), primary production can increase with consumers because they recycle nutrients that limit primary production [22], thus food webs are simultaneously “top down” and “bottom up” controlled. Consequently, most earth system and global biogeochemistry models include consumers, and sometimes consumers of consumers; however, there is little consensus on the configuration of food web structure or number of trophic levels to include, even though it significantly changes global scale primary production and plankton biogeography [23]. Developing models for microbial food webs is particularly challenging due to the extreme diversity of these communities [24], which cannot be explicitly represented in models due to our insufficient information on growth characteristics and predator-prey connectivity. As a consequence, modeled microbial food webs are aggregated and highly truncated, and “mortality closure” terms are introduced to approximate higher trophic levels not explicated represented in the model [25]. In the original work on food web network analysis, May [26] showed that large, randomly constructed food webs are unstable. Ever since that initial analysis, now referred to as May’s Paradox [27], there have been numerous publications that use stability as the central criterium for theoretical food web structure synthesis [e.g., 28]. Observations of microbial community dynamics, however, reveals that many appear unstable [29], while biogeochemistry remains near steady state. Therefore, an approach that differs from stability is needed to develop food web networks for models, especially for modeling microbial communities.

The objective of this work was to develop an MEP modeling approach to predict the food web structure of a heterotrophic microbial community that is defined by two palatability matrices, (bacteria), and **G** (consumers), where the former matrix describes which substrates each bacteria uses, and the latter matrix specifies which prey each consumer ingests. The elements of **B** and **G** are determined from an optimization problem in which entropy production is maximized over a specified interval to time based on the MEP principle. We investigated two optimization intervals, one short term (2 days; STO) and one long term (53 days; LTO). This modeling work was developed as part of a larger project that uses stable carbon isotope labels to experimentally track the flow of five substrates (glucose, xylose, acetate, ethanol and methanol) into bacteria and their predatory protists in chemostat-based bioreactors. Only model development and simulations will be presented here, and while the approach is general, the model structure is based on modeling the experimental setup.

## Methods

### MEP for non-steady-state systems

The current derivation of MEP only applies to steady-state systems, and no theory has yet been developed for transient processes, but some ideas have been proposed [30, 31]. Since most systems of interest are seldom at steady state, our initial work focused on extending the theory using a combination of microcosm experiments coupled with numerical studies that focused on microbial communities [19]. This work led to the conjecture that abiotic systems maximize instantaneous entropy production. That is, they take the steepest decent trajectory down a free energy surface, such as the path taken by a rock rolling down a hill that dissipates gravitational potential energy by converting it into heat. But the steepest decent trajectory may not produce the greatest amount of entropy when integrated over time. For instance, the rock may get stuck halfway down the hill in a ditch. In contrast, we have proposed [15, 20] that livings systems maximize entropy production averaged over time, which allows them to traverse alternate paths on a free energy surface that avoids barriers, such as the ditch halfway down the hill. To achieve this, living systems employ temporal strategies using information acquired by evolution and culled by natural selection, which is stored in their genomes. In fact, temporal strategies are hallmarks of biology, such as circadian rhythms [32], resource storage [33, 34], anticipatory control [35, 36], dormancy [37, 38], organism life cycles [39] and likely others we have yet to discover. However, temporal strategies are seldom incorporated in food web or biogeochemical models and our approach attempts to rectify this oversight.

Continuing the rock-hill analogy, the biological solution would have the rock take a path to the left or right of the steepest descent path, so the instantaneous entropy produced would be lower than the abiotic path; however, by avoiding the ditch, the rock could proceed to the bottom of the hill, thereby fully dissipating the gravitational potential and producing more entropy. We note, the entropy being referred to in MEP, and the rock-hill analogy, is not the state entropy, *S*, which can either increase or decrease in an open system (for instance, the entropy of a refrigerator decreases when turned on), but rather the entropy produced within a system caused by irreversible processes, denoted 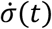 that has units of J K^-1^ d^-1^, and is always equal to or greater than zero as required by the Second Law [17, 40]. For the rock-hill case, it’s the conversion of gravitational potential energy into heat. We can formalize the above statements regarding the distinction between abiotic versus biotic processes as follows,

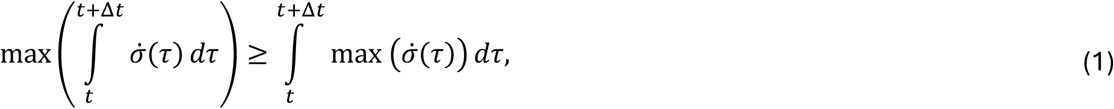

where Δ*t* is an interval of time that biology has evolved to operate over, such as one day for circadian rhythms, but much longer time intervals can exist, such as storing fat to survive food shortages or an organism’s full life cycle. The left side of Eq. (1) represents biology and the temporal strategies it implements that attempt to maximize 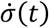 integrated over Δ*t*, while the right side captures abiotic processes that take the steepest decent trajectory, like a rock rolling down a hill, that maximize instantaneous entropy production, max (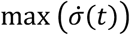. When Eq. (1) is true, biology has an advantage over abiotic processes in energy dissipation and it will likely dominate entropy production, but if an abiotic process can dissipate more free energy than biology over Δ*t*, then the abiotic process will be the preferred pathway for energy dissipation, biology will be excluded, and Eq. (1) will be false. While an abiotic system may attain much higher values of 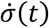 at certain instances, like a flash of fire, the area under the 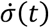 versus time curve will be lower than what a biologically catalyzed system can achieve when Eq. (1) is true.

### Bacteria and consumer growth equations, growth efficiency, and entropy production

To implement Eq. (1) for microbial communities, we represent bacterial growth on substrate *s*_*j*_ with two reactions, one representing substrate going to biomass *b*_*i*_ (so called anabolic reaction) and the other representing the combustion of the substrate to CO_2_ and water (catabolic reaction), then combine the two reactions with a growth efficiency variable, 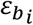 to give an overall reaction for bacterial growth that has this general form,

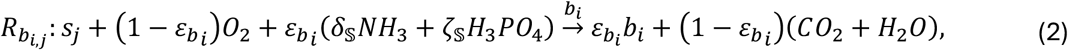

where *δ*_𝕤_ and *ζ*_𝕤_ are from the elemental composition of biomass given by 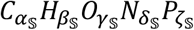, and the nomenclature 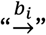 signifies the reaction is catalyzed by *b*_*i*_ (i.e., the reaction rate depends on the concentration of *b*_*i*_ and is also autocatalytic in this case). The substrate, *s*_*j*_, has a similarly defined composition, 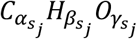, but lacks N and P to match the substrates used in the actual experiment. For clarity, we have omitted the stoichiometric coefficients of Eq. (2) (see Supplemental material, Eq. (S4), for details) so that we can focus on the main aspect of reaction 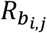. The stoichiometry and thermodynamics of 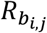 depends on 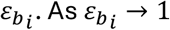, nearly all the substrate goes to produce biomass, and the reaction free energy, 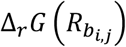, is near zero. At the other extreme, as 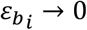, all the substrate is combusted to CO_2_ and water, resulting in no biomass synthesis, but a considerable amount of entropy is produced, as the chemical potential energy is just converted to heat (also see Eq. (S5)). In fact, we can easily calculate the entropy production from reaction 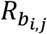, which is given by,

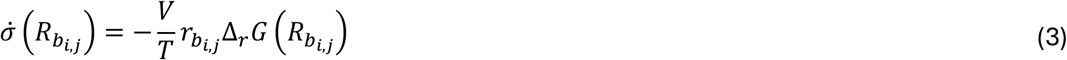

where *V* and *T* are the volume and temperature of the system, respectively, and 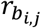 is the rate of reaction 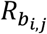. The reaction rate depends on a substrate uptake rate expression, the bacterial preference for substates (i.e., the **B** matrix, see below) and the bacteria concentration. We use an adaptive Monod equation [19, 41] for the substrate uptake rate where the classic Monod parameters (*μ*^*Max*^ and *k*_*M*_) depend non-linearly on the value of 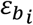, as well as a thermodynamic driver [42] that depends on 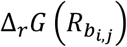, so that the reaction rate goes to 0 as 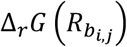 approaches 0 (i.e., equilibrium). The adaptive Monod equation captures growth strategies from oligotrophy, when 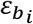 is closer to 0, to copiotrophy, when 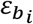 is closer to 1. The growth model for consumers eating bacteria, or other consumers, is nearly identical to that described for bacterial growth on substrates, except the “substrate” is a bacterium, another consumer, or itself, as defined by the **G** matrix, and consumers remineralize N and P as ammonium (NH_3_) and inorganic phosphate (H_3_PO_4_) (see Eqs. (S8), (S12), and (S14) and below).

### Food web structure

The food web structure is defined by the matrices **B** and **G** that determine the preference of each bacteria “species”, or instance, *b*_*i*_, for the five substrates used in the experiment, and the preference of each consumer for prey, respectively (Fig. 1). The **B** matrix has the dimensions of *n*_*b*_ × *n*_*s*_ and that of the **G** matrix is *n*_*g*_ × (*n*_*b*_ + *n*_*g*_), where *n*_*s*_, *n*_*b*_, and *n*_*g*_ are the number of substrates, bacteria, and consumers, respectively. For the simulations run for this study, *n*_*s*_, *n*_*b*_, and *n*_*g*_ were set to 5, 10, and 5, respectively, where the substrates used in the experiment and simulations were glucose, xylose, acetate, ethanol, and methanol. We did not consider more complex networks that could include cross-feeding, bacterial predators or pathogens, nor consumer mixotrophs, which would significantly increase the number of unknowns in the model (see full transfer matrix, Fig. S1). Because consumers can prey on other consumers and themselves, the **G** matrix also determines the number of trophic levels in the network (Fig. 1) and the extent of cannibalism.

**Fig. 1.**
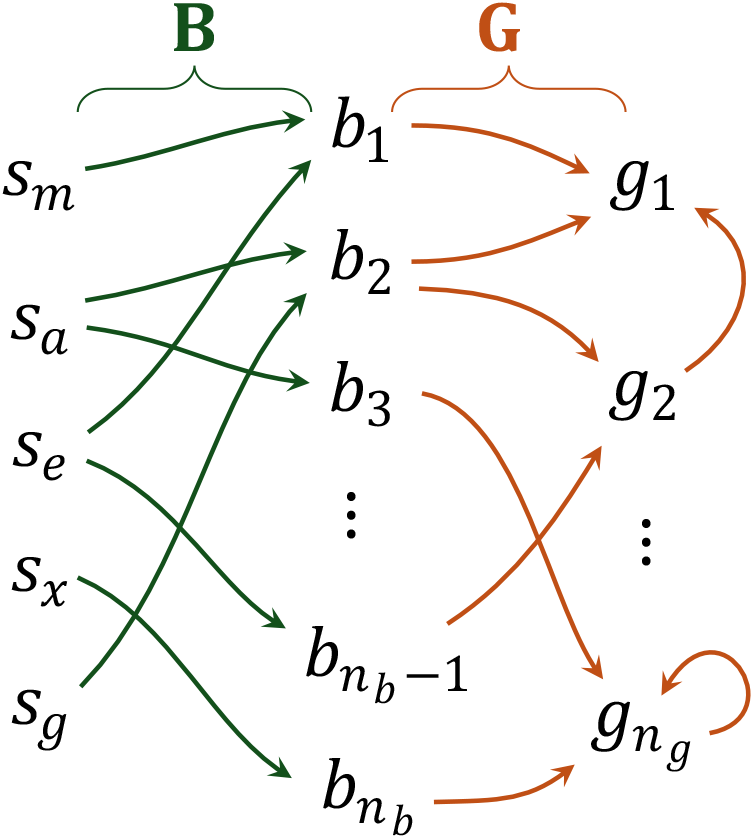
Example of how the **B** matrix defines which substrate (*s*_*m*_: methanol, *s*_*a*_: acetate, *s*_*e*_: ethanol, *s*_*x*_: xylose, *s*_*g*_: glucose) are used by the *n*_*b*_ bacteria, *b*_*i*_, and the **G** matrix defines which bacteria, or consumers, each of the *n*_*g*_ consumers, *g*_*i*_ ingests. Because a consumer can eat other consumers, including itself (i.e., cannibalism), the number of trophic levels is also set by **G**.

A set of ordinary differential equations derived from mass balances and biologically catalyzed reaction rates (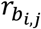 and 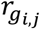) describe the dynamics between carbon substrates, bacteria, and consumers in a chemostat of volume *V* that is being fed a sterile minimal salts medium with the five carbon sources at a volumetric flow rate of *F*_*L*_ (Eqs. (S15-S17)). The chemostats are sparged with air at a gas volumetric flow rate of *F*_*G*_, and a similar set of mass-balance derived differential equations track the concentration of dissolved oxygen and inorganic carbon (i.e., carbonate species) in the medium, as well as the partial pressures of oxygen and carbon dioxide in the bioreactors headspace (Eqs. (S18-S22)). Since the experimental medium was designed to be inorganic phosphate (H_3_PO_4_) limited, bacterial reaction rates, 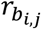, are also limited by H_3_PO_4_ concentration (Eq. (S11)), which is also included in the set of governing differential equations (Eq. (S22)). Consumers also remineralize H_3_PO_4_ as part of their predation activities (Eq. (S8)). For simulations presented in Results, the chemostat dilution rate (*F*_*L*_ /*V*_*L*_) was set to 0.1 d^-1^ (0. *L d*^™1^/3.0 *L*), and the concentration of the 5 substrates were set to (mmol m^-3^): glucose, 50; acetate, 167.5; ethanol, 108.3; methanol, 206.1; xylose, 59.69. The substrate concentrations were designed so that each provides the same free energy input of 147 J per L of medium. Other parameter values can be found in the *.inp files included in the supplementary material and a GitHub repo [43].

### MEP optimization

The only variables in the growth models for bacteria and consumers are 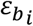 and 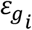, which set the stoichiometries, the thermodynamics, and the rates of reactions for 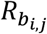 and 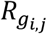, as well as the entropy production associated with growth and respiration. The other unknowns are the elements of the **B** and **G** matrices that define food web structure; consequently, there are a total of 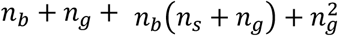 unknown parameters, *n*_*u*_, that need to be set to define the model. For our simulations here, where *n*_*s*_ = 5, *n*_*b*_ = 10, and *n*_*g*_ = 5, the total number of unknowns is *n*_*u*_ = 140. In conventional food web or biogeochemistry models, these parameters would be set via a combination of literature data and guesses, assumptions and theories on matrix structure, and data assimilation, assuming data were available. However, under the conjecture that systems organize to maximize entropy production, Eqn (1), we can define an optimization problem that determines the *n*_*u*_ parameter values by maximizing entropy production. Since this is a biological system where Eq. (1) applies, entropy production is maximized over an interval of time, *δ*_*MEP*_, as defined by Eqs. (S25, S26), but the appropriate choice of *δ*_*MEP*_ is unknown, as it reflects the forecasting time scale over which biology has evolved to operate. However, the variational approach does reduce the problem from 140 unknowns down to 1, which we can manipulate to understand how *δ*_*MEP*_ influences the solution.

### Food web metrics

In addition to model output for the concentration of substrates, bacteria, grazers, and environmental variables (Eqs. (S15-S22)), we also calculate two metrics of the food web structure for different values of *δ*_*MEP*_. The substrate diversity of *b*_*i*_bacteria, that is, the effective number of substrates *b*_*i*_ bacteria use, was based on Hill numbers [44] of order 2, ^*q*^*D*_*i*_ (*q* =, or arithmetic mean weighting) calculated from the row vectors of the **B** matrix (Eq. (S27)). For instance, if the row of the **B** matrix for *b*_*i*_ bacteria is [0.9 0.0 0.0 0.0 0.0], then ^2^*D*_*i*_ equals 1.23, reflecting the fact that the bacteria places most of its effort into consuming substrate 1, although it will weakly consume the others if they are present. The other metric, calculated from the **G** matrix, was the tropic position, 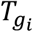, of consumer *g*_*i*_ (Eq. (S28)), where 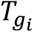 was derived from the full transfer matrix (Fig. S1) as described by Levin [45]. Because foods webs can include cycles, such as consumers 1 and 2 eat each other, as well as any cannibalism, the number of trophic positions is effectively unbounded and depends on the value of 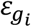 that sets the number of passes around a cycle. Note, in simulations where *b*_*i*_ or *g*_*i*_ never exceeded their initial values (i.e., no growth occurred), the corresponding values for ^2^*D*_*i*_ or 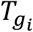 were set to 0, respectively. All source code for the model runs is available as a GitHub repo [43].

## Results

To examine how the time interval over which integrated entropy production is maximized effects the solution and the food web structure of the microbial community, we present two model runs in which *δ*_*MEP*_ was set to either 2 or 53 days. Both simulations were run for 53 days to reflect the duration of the actual experiment; however, the 140 combined elements in the two vectors, *ε*_*b*_ and *ε*_*g*_, and two matrices, **B** and **G**, were optimized using hyperBOB [46] to maximize entropy production only over the initial time interval [0, *δ*_*MEP*_]. For instance, when *δ*_*MEP*_ = d, the elements in the vectors and matrices were adjusted so that the integrated entropy production was maximized over the first 2 days of simulation. Once the optimal parameters were found over [0, *δ*_*MEP*_], the model integration continued from *δ*_*MEP*_ to *t*_*f*_, where *t*_*f*_ = 5 d. Because these types of optimization problems are typically replete with local optima, the optimization was run in parallel on 200 CPUs and only the best solutions are presented here; however, all solutions were used for the two food web metrics, substrate diversity, ^2^*D*_*i*_ and trophic position, 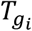 (Eqs. (S27) and (S28)).

### Short versus long-term optimization intervals

Below, we will use Short-Term Optimization (STO) to refer to the simulation where *δ*_*MEP*_ was set to 2 days, and Long-Term Optimization (LTO) for the case where *δ*_*MEP*_ equaled 53 days. Instantaneous entropy production (EP),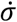, and integrated (i.e., cumulative) entropy produced, *σ*, for the two simulations (Fig. 2) shows that integrated EP over the first 2 days was greater for the STO than the LTO simulation (Fig. 2a, solid vs dashed red lines), and the highest EP rate over the entire simulation of 2.36 J K^-1^ d^-1^ occurred at 1.73 days for the STO case (Fig. 2a, solid vs dashed black lines). While the peak instantaneous EP was lower in the LTO simulation, the integrated EP rate exceeded that of the STO beginning at 2.58 days, and by the end of the simulation, the LTO solution generated 41.5 J K^-1^ of entropy, while the STO only produced 23.1 J K^-1^ (Fig. 2b, red lines). After approximately 15 days, both solutions attain steady-state operation with constant EP rate of 0.39 and 0.70 J K^-1^ d^-1^ for the STO and LTO, respectively.

**Fig. 2.**
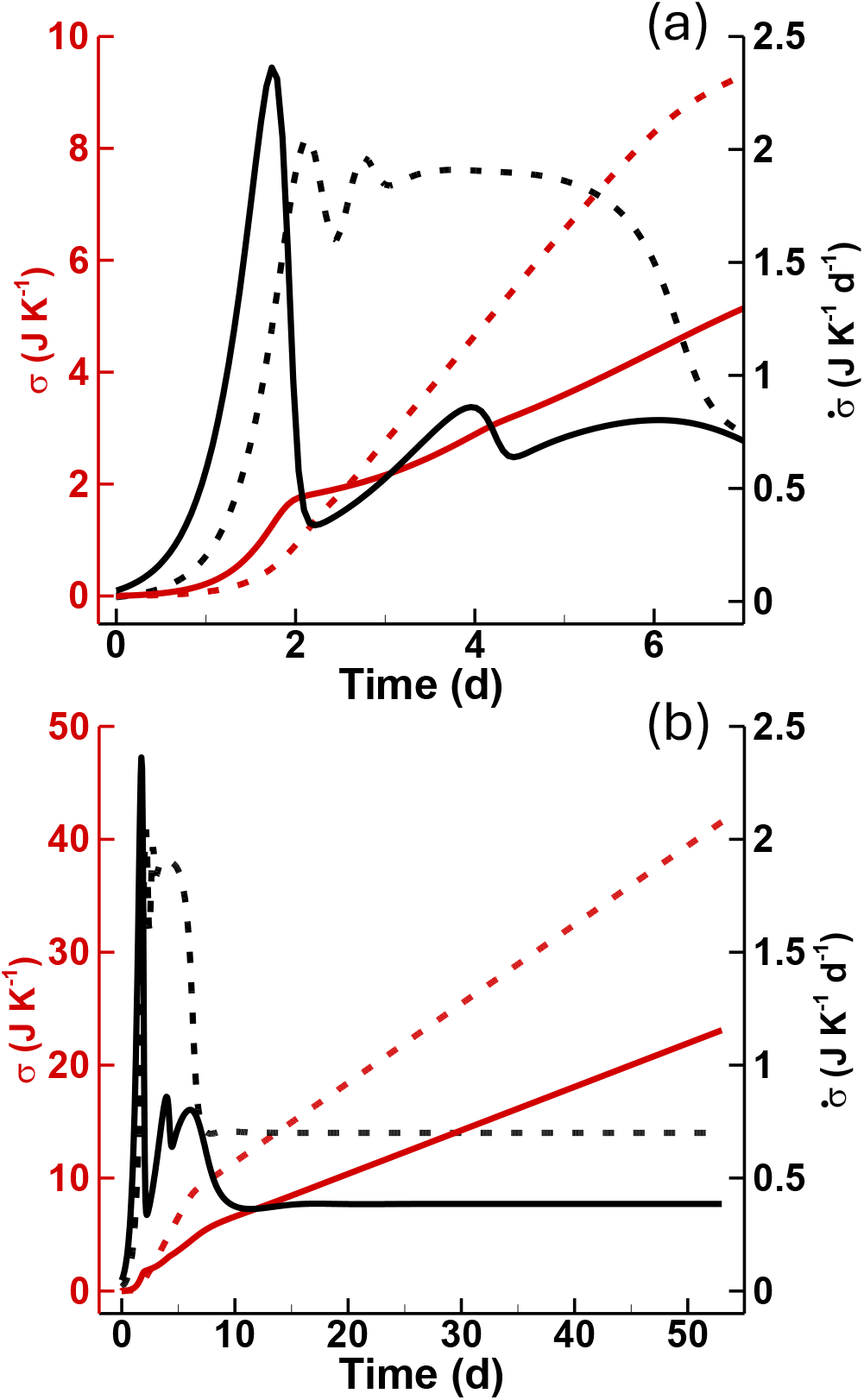
Instantaneous entropy production (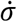, black lines) and integrated (cumulative) entropy generated (*σ*, red lines) from irreversible processes when the latter is maximized over [0, 2] days (*δ*_*MEP*_ =, STO solid lines) versus maximized over [0, 53] days (*δ*_*MEP*_ = 5, LTO dashed lines) plotted for (a) the first 7 days and (b) all 53 days of the simulation.

To attain maximum EP over 2 days, the STO solution selected for all but one bacteria instance to have similar rapid growth kinetics that peaked in biomass at ∼ 9.2 µM around day 1, with *b*_6_ bacteria attaining a secondary peak of 25 µM at 6.9 days (Fig. 3a). After day 15, all bacteria exhibited coexistence and steady-state concentrations, except *b*_7_ bacteria that were washed out of the bioreactor at the start. Coexistence occurred because all 9 bacteria had similar growth efficiencies of approximately 0.2, while the optimization assigned *b*_7_ bacteria an efficiency of ∼1 that prevented their growth (Fig. S2a). The LTO solution exhibited significantly different dynamics that involved only 4 viable bacteria that displayed a pattern of succession (Fig. 3b). The *b*_2_ bacteria dominated first, but those bacteria were quickly replaced by *b*_1_ bacteria, which in turn were replaced by *b*_5_ bacteria. By day 10, the community was dominated by *b*_4_ bacteria, with all the other bacteria displaced in a classic competitive exclusion scenario. Bacteria growth efficiencies for *b*_2_, *b*_1_, *b*_5_ and *b*_4_, followed a sequential decline in values from 0.232, 0.196, 0.191, to 0.142, respectively (Fig. S2a).

**Fig. 3.**
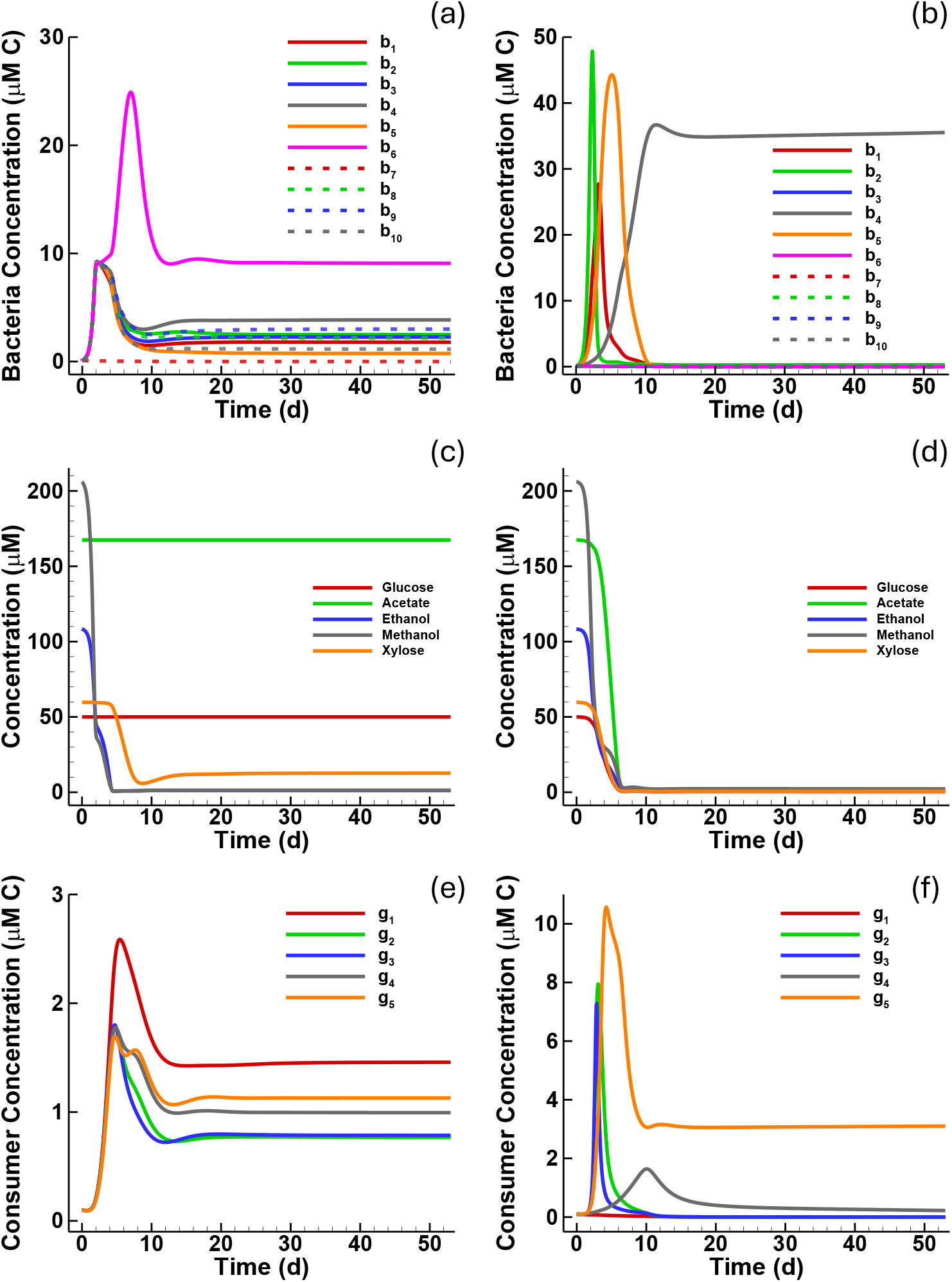
Predicted concentrations (μM) of (a,b) bacteria, (c, d) substrates, and (e, f) consumers when integrated entropy production is maximized, *δ*_*MEP*_, over (left: a, c, e) STO 2 days versus (right: b, d, f) LTO 53 days.

The STO solution resulted in rapid bacterial uptake of methanol and ethanol and only a very slight uptake of xylose in the first two days of the simulation, which defines the STO interval (Fig. 3c). Rapid uptake of xylose then followed over days 4.3 to 7.8. Neither glucose nor acetate were consumed during the entire STO simulation. This pattern of substrate preference in the STO solution is evident in the **B** matrix, as only those columns that correspond to ethanol and methanol exhibit high values (Fig. S3a). While the **B** matrix did show that *b*_7_ bacteria allocated significant resources to glucose, acetate, and xylose uptake, the STO solution prevented *b*_7_’s growth by assigning it a growth efficiency of 1. Only *b*_6_ bacteria consumed xylose in the STO (*B*_6,5_ = 0.0 8). In the LTO solution, all substrates were rapidly consumed and brought down to nearly 0 concentration (Fig. 3d), but the first bacteria to dominate, *b*_2_, had a preference for ethanol and methanol, followed by *b*_1_ bacteria that preferred glucose, ethanol and xylose, and *b*_5_ and *b*_4_ bacteria allocated most of their resources to glucose and xylose uptake, as based on the optimized values of the **B** matrix (Fig. S3b).

Consumers also exhibited different dynamics between the STO and LTO solutions. Like the bacteria in the STO solution, consumers from the STO all had similar bursts in growth, followed by a steady-state coexistence after ca. 20 days (Fig. 3e), and all had similar growth efficiencies of approximately 0.2 (Fig. S2b). Most consumers dined on 2 or 3 bacteria, and 3 or 4 consumers (including some cannibalism), except *g*_5_ consumer that only ingested *b*_5_ bacteria (Fig. S4a). Note, column 7 of **G** for the STO solution (Fig. S4a) can be ignored since *b*_7_ bacteria were washed out of the bioreactor and the prey preference matrix is weighted by prey abundance (Eq. (S13)). Consumers in the LTO solution (Fig. 3f) exhibited, like the bacteria, competitive exclusion, with *g*_5_ consumer dominating by the end of the simulation, while *g*_1_ consumer did not grow due to its growth efficiency of 1 (Fig. S2b). The grow efficiencies of the 4 viable consumers varied from 0.12 (*g*_4_) to 0.28 (*g*_3_), and the dominant consumer, *g*_5_, had an efficiency of 0.22. The consumer prey-preference matrix, **G**, was similar to that from the STO solution, but consumers only ingested 1 or 2 bacteria or 1 to 3 consumers (Fig. S4b). This decrease compared to the STO solution was due to fewer viable bacteria and consumers (only 4 of each) in the LTO solution. However, this lower trophic complexity was not evident in other solutions that performed nearly as well in terms of entropy production as the best solution.

#### Substrate diversity and trophic position metrics

Substrate diversity of the bacteria, ^2^*D*_*i*_, calculated from the **B** matrix (Eq. (S27)), and trophic position of the consumers, *T*_*gi*_, calculated from the **G** matrix (Eq. (S28)), were determined for all 200 local optima solutions produced for each of the STO and LTO simulations, which were ranked from 1 (best) to 200 (worst) (Fig. 4). The substrate diversity metric shows a clear difference between the STO and LTO solutions, where the diversity of substrates bacteria used in the LTO solutions were approximately 4, while those for the STO solutions were constrained between 1 and 2 substrates (Fig. 4a). The trophic position of the consumers was approximately 2.5 for the consumers in the STO solution, but *T*_*gi*_ varied significantly for consumers in the LTO solution, ranging from around 3 up to nearly 10 (Fig. 4b).

**Fig. 4.**
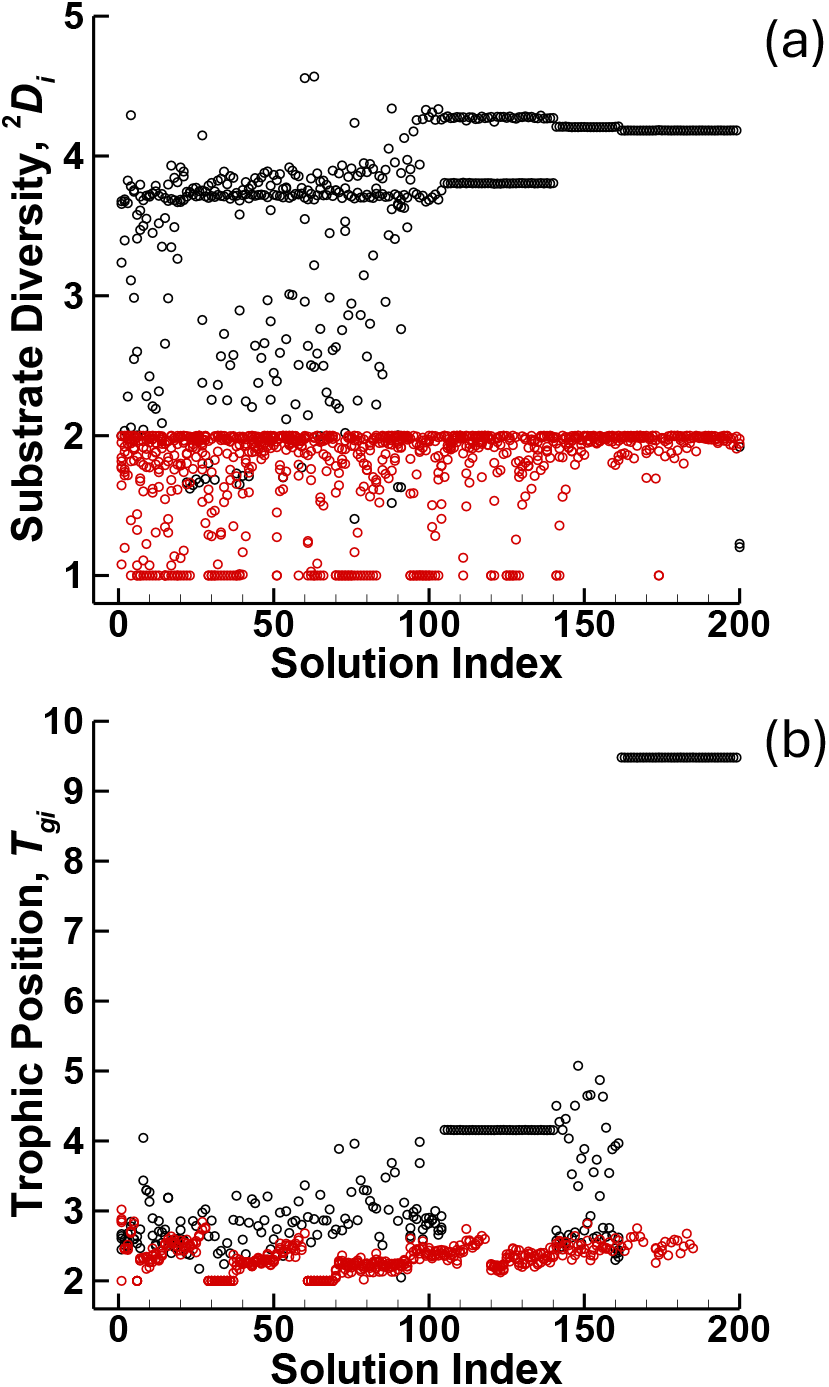
(a) Substrate diversity, ^2^*D*_*i*_, of the bacteria and (b) trophic position, *T*_*gi*_, of the consumers for the short-term optimization (STO, red circles) and long-term optimization (LTO, black circles) simulations that each located 200 local optima. Solution index 1 was the best optima found (and presented in other figures), while solution index 200 was the worst located. ^2^*D*_*i*_ and *T*_*gi*_ were not included here if *b*_*i*_ or *g*_*i*_ never exceeded their initial concentration values during the simulation.

## Discussion

The results from the short-term (STO) and long-term optimization (LTO) illustrate the importance of temporal strategies to how living systems can outcompete abiotic systems in terms of free energy dissipation. While the STO solution achieved the highest rate of entropy production and produced more entropy over the first 2 days of the simulations, the LTO solution maintained higher average entropy production. After 53 days, the LTO solution produced 1.8 times more entropy than the STO solution. The STO solution placed almost all available bacterial resources on methanol and ethanol consumption and oxidation, because methanol and ethanol’s free energies of combustion (−679.4 and −654.8 kJ (mol C)^-1^, respectively) are higher than those of glucose and acetate (−483.1 and - 425.5 kJ (mol C)^-1^, respectively) under the chemostat’s initial conditions. This resulted in high rates of entropy production for the STO, but glucose and acetate were unused, which explains why the substrate diversity index for all STO solutions was between 1 and 2 and the lower entropy production over 53 days compared to the LTO solution. The growth kinetics of 9 out of 10 bacteria in the STO solution were so similar in value that the STO solution ended with high bacterial diversity compared to the LTO solution that exhibited competitive exclusion with only one bacteria remaining at the end of the simulation. A small fraction of bacterial resources were placed on xylose consumption in the STO solution, which had the third highest free energy of combustion (−489.2 kJ (mol C)^-1^), but in general the STO solution selected for substrate specialists. Effectively, the STO solution operated closer to the steepest descent approach in energy dissipation, similar to the rock rolling down a hill. In contrast, the LTO solution allocated resources for both short- and long-term behavior that resulted in both rapid uptake, as well as complete utilization, of all substrates. To achieve this, the LTO solution selected for both opportunistic and gleaner bacteria [47], where the opportunistic bacteria with higher growth efficiencies, *b*_2_, *b*_1_, and *b*_5_ were able to grow rapidly at the start of the experiment when substrate concentrations were high, but were displaced by the gleaner bacteria with a lower growth efficiency, *b*_4_, once the substrate concentrations had been drawn down to low levels. The bacterial community succession in our LTO mimics natural bacterial succession in chemostats under steady-state conditions [29]. Overall, the bacteria in the LTO solution were substrate generalists. While this distribution of growth characteristics was unable to generate the highest instantaneous entropy production (i.e., it was not the steepest descent approach), over longer time scales it more effectively used the available resources and produced almost twice as much entropy as the STO solution.

The trophic structure of the STO and LTO solutions based on the trophic position metric, *T*_*gi*_, showed that the LTO solution exhibited more trophic levels and had a higher dependence on consumers than did the STO solution. For some of the lower ranked STO solutions (186-200), no viable consumers were selected, and when the optimization interval, *δ*_*MAP*_, was reduced to 1 day (data not included here), even the best STO solutions did not include any consumers. The consumers in the LTO solution were approximately 1 trophic position higher than in the STO solutions for the best ranked solutions. The LTO solution also produced consumers with *T*_*gi*_ values approaching 10 for the lower ranked solutions. For those solutions inspected, the high *T*_*gi*_ values were attained by a consumer placing preferences on a single dominate bacteria combined with a cannibalism loop, which increased *T*_*gi*_. The consumers provide two important functions with respect to entropy production: reducing the standing stock of biomass and recycling nutrients, such as phosphate. Because bacteria and consumers can be oxidized to CO_2_ and water, entropy production can be increased via their consumption if their metabolic contribution is no longer needed. For instance, in a batch system, once a substrate is completely consumed, those bacteria specialized for that substrate are no longer needed, so they should be combusted, which consumers accomplish. From the perspective of the MEP model, the objective is not to build biomass, as that does not produce entropy, but biomass is required to catalyze the destruction of chemical potential energy that does produce entropy. The model objective is to build just enough catalytic biomass to dissipate the available chemical free energy, and consumers are a means to achieve a constraint on biomass. Similarly, consumers facilitate the reallocation of community resources to other community metabolic functions, such as switching community metabolism from methanol to glucose by changing the community composition through directed bacterial predation. Of course, consumers also recycle nutrients trapped in biomass that are needed by the primary producers (bacteria here) for growth, so the presence of consumers, in the words of Lotka [48], ‘accelerate the circulation of matter through the life cycle, both by “enlarging the wheel,” and by causing it to “spin faster.”‘, which results in higher entropy production [17]. However, the development of consumer biomass takes time, so under the STO, they are not as important, or not needed at all, if the time scale is sufficiently short.

The objective of this study was to develop an approach to generate food web structures for microbial communities (i.e., the **B** and **G** connectivity matrices). There is a considerable body of theoretical research that concerns constructing synthetic connectivity matrices for understanding classic macroscopic food webs, such as the Cascade Model [49], the Niche Model [50], the Phylogenetic Niche Model [51] and the Preferential Preying Model that underlies May’s Paradox [27]. These approaches are often based on the heuristic that unstable dynamics should be culled from the solution space, such as in qualitative network analysis [52], because collective wisdom dictates that complex natural communities should be stable [53]. However, microbial systems contain more than 10^9^ organisms L^-1^ or 10^12^ organisms kg^-1^ in aquatic or terrestrial soils, respectively [54], that disperse readily, and small volumes can contain upwards of 10,000 species or more, each with different growth kinetics [24, 55]. For microbial systems, stability may not be a defining feature, and there is experimental support for this concept. For instance, we found unstable bacterial communities underlaid near constant methane oxidation rates [19, 56]. Similarly, Graham et al. [57] observed fairly stable nitrification rates that were supported by unstable community dynamics, Fernández et al. [58] found an unstable methanogenic community but stable methanogenesis, and Benincà et al. [59] observed chaotic plankton communities in long term experiments as well. While stability may be a compelling objective for food web analysis, it may be the wrong solution from an energy dissipation perspective, at least for microbial ecosystems, where May’s [26] conclusions may not be paradoxical. Our MEP-based approach does not require assumptions on stability to predict microbial food web structures, but it needs to be tested experimentally, which we are currently pursing [60].

## Supporting information

Governing Equations, Figures and parameters

## Acknowledgments

This research was funded by the Simons Foundation grant 549941FY22 (J.J.V., O.M.A.) and NSF awards: 1655552 (J.J.V., O.M.A., J.A.H.), 2224608 (J.J.V.)

