## Supplementary material for "Deriving microbial food web structure by maximizing entropy production over variable timescales": Governing Equations, Figures and parameters

#### Bacteria Reactions

Growth of bacteria on substrates is represented by two chemical reactions, one capturing catabolism (energy acquisition) per unit C, given by,

$$R^C_{b_{i,j}}:\quad \frac{1}{\alpha_{s_j}} s_j + a^C_{s_j} O_2 \overset{b_i}{\rightarrow} H_2CO_3 + b^C_{s_j}\mathrm{H_2O}, \tag{S1}$$

and the other representing anabolism (growth) per unit C, given by,

$$R^A_{b_{i,j}}:\quad \frac{1}{\alpha_{s_j}} s_j + \frac{1-a^A_{s_j}}{\alpha_{\mathbb{S}}}(\delta_{\mathbb{S}} NH_3 + \zeta_{\mathbb{S}} H_3PO_4) \overset{b_i}{\rightarrow} \frac{1-a^A_{s_j}}{\alpha_{\mathbb{S}}} b_i + a^A_{s_j} H_2CO_3 + b^A_{s_j}\mathrm{H_2O}, \tag{S2}$$

where $s_j$ is one of $n_s$ substrates that matches the five used in the stable isotope probing (SIP) experiment (methanol, acetate, ethanol, xylose or glucose) with elemental composition given by $C_{\alpha_{s_j}} H_{\beta_{s_j}} O_{\gamma_{s_j}}$, $b_i$ is bacteria $i$, of which there are $n_b$ instances that all have the same elemental composition given by $C_{\alpha_{\mathbb{S}}} H_{\beta_{\mathbb{S}}} O_{\gamma_{\mathbb{S}}} N_{\delta_{\mathbb{S}}} P_{\zeta_{\mathbb{S}}}$, and $r^C_{b_{i,j}}$ and $r^A_{b_{i,j}}$ designate catabolic and anabolic reaction rates catalyzed by bacteria $b_i$ utilizing substrate $s_j$, based on reactions $R^C_{b_{i,j}}$ and $R^A_{b_{i,j}}$respectively. The four stoichiometric coefficients, $a^C_{s_j}$, $b^C_{s_j}$, $a^A_{s_j}$ and $b^A_{s_j}$, are readily determined from elemental balances around H and O in each reaction. For instance, $a^A_{s_j}$ is given by,

$$a^A_{s_j} = \frac{\alpha_{s_j}(\beta_{\mathbb{S}} - 2\gamma_{\mathbb{S}} - 3\delta_{\mathbb{S}} + 5\zeta_{\mathbb{S}}) - \alpha_{\mathbb{S}}\left(\beta_{s_j} - 2\gamma_{s_j}\right)}{\alpha_{s_j}(\alpha_{\mathbb{S}} + \beta_{\mathbb{S}} - 2\gamma_{\mathbb{S}} - 3\delta_{\mathbb{S}} + 5\zeta_{\mathbb{S}})}. \tag{S3}$$

The overall reaction representing grown of each bacteria on the $n_s$ substrates is constructed by multiplying $R^A_{b_{i,j}}$ by $\varepsilon_{b_i}$ and adding it to $R^C_{b_{i,j}}$ multiplied by $(1-\varepsilon_{b_i})$, where $\varepsilon_{b_i}$ is the thermodynamic or growth efficiency parameter, bounded between 0 and 1, that is determined by maximizing entropy production over a specified interval of time. The overall chemical reaction describing bacterial growth is then,

$$R_{b_{i,j}}:\quad \frac{1}{\alpha_{s_j}} s_j + (1-\varepsilon_{b_i}) a_{s_j}^C O_2 + \varepsilon_{b_i} \frac{1-a_{s_j}^A}{\alpha_{\mathbb{S}}} (\delta_{\mathbb{S}} NH_3 + \zeta_{\mathbb{S}} H_3PO_4) \xrightarrow{b_i} \varepsilon_{b_i} \frac{1-a_{s_j}^A}{\alpha_{\mathbb{S}}} b_i + \left(\varepsilon_{b_i} a_{s_j}^A + 1 - \varepsilon_{b_i}\right) H_2CO_3 + \left(\varepsilon_{b_i} b_{s_j}^A + (1-\varepsilon_{b_i}) b_{s_j}^C\right) \mathrm{H_2O}, \tag{S4}$$

where the Gibbs free energy of reaction for the catabolic, $\Delta_r G(R_{b_{i,j}}^C)$, and anabolic, $\Delta_r G(R_{b_{i,j}}^A)$, reactions $R_{b_{i,j}}^C$ and $R_{b_{i,j}}^A$ are calculated from Gibbs free energies of formation obtained from Alberty [1] at the given temperature, pH, and ionic strength, and the concentrations of all chemical species via the natural logarithm of the reaction quotient, $Q$, temperature, and the ideal gas constant (i.e., $\Delta_r G = \Delta_r G^\circ + RT \ln(Q)$). The overall reaction free energy for Eq. (S4) is then given by,

$$\Delta_r G\left(R_{b_{i,j}}\right) = \varepsilon_{b_i} \Delta_r G\left(R_{b_{i,j}}^A\right) + (1-\varepsilon_{b_i}) \Delta_r G\left(R_{b_{i,j}}^C\right), \tag{S5}$$

which is strongly controlled by the value of $\varepsilon_{b_i}$. As $\varepsilon_{b_i} \to 0$, the substrate is completely oxidized, and the free energy (chemical potential energy) is dissipated as heat, resulting in maximum instantaneous entropy production; as $\varepsilon_{b_i} \to 1$, chemical potential energy is largely conserved, resulting in little to no entropy production (see below).

### Consumer Reactions

The reactions of the consumers are handled similarly to that for bacteria. The catabolic and anabolic reactions representing the growth of consumers, $g_i$, (i.e., protistan grazers) on prey, $p_j$, that includes both bacteria and the consumers themselves (i.e., $p_j = \begin{bmatrix} b_j \\ g_j \end{bmatrix}$ and Fig. 1), on a unit carbon bases are given by,

$$R_{g_{i,j}}^{C}: \quad \frac{1}{\alpha_{\mathbb{S}}} p_j + a_{p_j}^{C} O_2 \xrightarrow{g_i} H_2CO_3 + b_{p_j}^{C} \mathrm{H_2O} + \frac{\delta_{\mathbb{S}}}{\alpha_{\mathbb{S}}} NH_3 + \frac{\zeta_{\mathbb{S}}}{\alpha_{\mathbb{S}}} H_3PO_4 \tag{S6}$$

$$R_{g_{i,j}}^{A}: \quad \frac{1}{\alpha_{\mathbb{S}}} p_j \xrightarrow{g_i} \frac{1}{\alpha_{\mathbb{S}}} g_i, \tag{S7}$$

where we have made the simplification that the elemental composition of the consumers, $g_i$, is the same as bacteria, $b_i$, and is also given by $C_{\alpha_{\mathbb{S}}} H_{\beta_{\mathbb{S}}} O_{\gamma_{\mathbb{S}}} N_{\delta_{\mathbb{S}}} P_{\zeta_{\mathbb{S}}}$. While this simplification is not necessary for the model, we do not have information regarding compositional differences between prey and consumer, so we have chosen the mathematically cleaner version. The two consumer reactions are also combined via the thermodynamic efficiency parameter, $\varepsilon_{g_i}$, to produce the overall reaction for growth and respiration of consumers,

$$\begin{aligned} R_{g_{i,j}}: \quad \frac{1}{\alpha_{\mathbb{S}}} p_j + (1 - \varepsilon_{g_i}) a_{p_j}^{C} O_2 &\xrightarrow{g_i} \frac{\varepsilon_{g_i}}{\alpha_{\mathbb{S}}} g_i \\ &+ (1 - \varepsilon_{g_i}) \left( H_2CO_3 + b_{p_j}^{C} \mathrm{H_2O} + \frac{\delta_{\mathbb{S}}}{\alpha_{\mathbb{S}}} NH_3 + \frac{\zeta_{\mathbb{S}}}{\alpha_{\mathbb{S}}} H_3PO_4 \right). \end{aligned} \tag{S8}$$

The equations for Gibbs free energy of reaction, $\Delta_r G \left( R_{g_{i,j}} \right)$, are similarly derived to those described above for bacteria.

### Reaction Rates

Reaction rates for all bacteria, $r_{b_{i,j}} \in \mathbb{R}^{n_b \times n_s}$**,** take the general form of,

$$r_{b_{i,j}} = \Phi_{b_{i,j}} B'_{i,j} c_{b_i}, \tag{S9}$$

where the matrix $\mathbf{\Phi}_b \in \mathbb{R}^{n_b \times n_s}$ is the substrate specific-uptake rate based on an adaptive Monod model [2] (see Eq. (S11) below), and $\mathbf{B}'$ is a substrate weighted $\mathbf{B} \in \mathbb{R}^{n_b \times n_s}$ matrix. The $\mathbf{B}'$ matrix is given by,

$$\mathbf{B}'_i = \frac{\mathbf{B}_i \odot \mathbf{c}_s}{\mathbf{B}_i \cdot \mathbf{c}_s}, \tag{S10}$$

where $\mathbf{B}_i \in \mathbb{R}^{1\times n_s}$ is the $i^{th}$ row vector of $\mathbf{B}$ and analogously for $\mathbf{B}'_i$, and $\odot$ and $\cdot$ are the elementwise (Hadamard) and dot products, respectively. Rows of the substrate preference matrix, $\mathbf{B}_i$, sum to unity, and determine how much catalyst from bacteria $i$ is allocated to each of the $n_s$ substrates. For example, when $n_s = 5$, bacterial specialists can be represented with all the weight on one substrate, such as $\mathbf{B}_i = [0\ 1\ 0\ 0\ 0]$, while generalists spread metabolic resources across the 5 substrates, such as $\mathbf{B}_i = [0.2\ 0.3\ 0.2\ 0.1\ 0.2]$.

The substate specific-uptake rate by bacteria $i$ using $s_j$, $O_2$, $NH_3$ and $H_3PO_4$ is given by,

$$\Phi_{b_{i,j}} = \nu^* \varepsilon_{b_i}^2 \left(\frac{c_{s_j}}{c_{s_j} + \frac{1}{\alpha_{s_j}}\kappa^* \varepsilon_{b_i}^4}\right)\left(\frac{c_{O_2}}{c_{O_2} + a_{s_j}^C \kappa^* \varepsilon_{b_i}^4}\right) \times \left(\frac{c_{NH_3}}{c_{NH_3} + \frac{1}{\alpha_{\mathbb{S}}}\delta_{\mathbb{S}}\left(1 - a_{s_j}^A\right)\kappa^* \varepsilon_{b_i}^4}\right)$$
$$\times \left(\frac{c_{H_3PO_4}}{c_{H_3PO_4} + \frac{1}{\alpha_{\mathbb{S}}}\zeta_{\mathbb{S}}\left(1 - a_{s_j}^A\right)\kappa^* \varepsilon_{b_i}^4}\right) F_T\left(\Delta_r G\left(R_{b_{i,j}}\right), n_{b_{i,j}}^e\right), \quad \text{(S11)}$$

where $F_T$ is the thermodynamic driver described by LaRowe [3] that inhibits the reaction as $\Delta_r G\left(R_{b_{i,j}}\right) / n_{b_{i,j}}^e \to 0$, $n_{b_{i,j}}^e$ is the number of electrons transferred in the catabolic reaction, given by $4a_{s_j}^C$, and $\nu^*$ and $\kappa^*$ are fixed constants independent of the substrate or phyla, and equal 350. $d^{-1}$ and 5000. mmol $m^{-3}$, respectively.

Reaction rates for consumers follow a similar form to that for bacteria, given by,

$$r_{g_{i,j}} = \Phi_{g_{i,j}} G'_{i,j} c_{g_i}, \quad \text{(S12)}$$

where $\mathbf{\Phi}_g \in \mathbb{R}^{n_g\times(n_b+\boldsymbol{n_g})}$ is the prey specific-uptake rate, $\mathbf{c}_g \in \mathbb{R}^{(n_b+n_g)\times 1}$ is the consumer concentrations, and $\mathbf{G}'$ is a prey-weighted $\mathbf{G}$ matrix which determines which prey each consumer eats. The prey-weighted $\mathbf{G}'$ matrix is given by,

$$\mathbf{G}_i' = \frac{\mathbf{G}_i \odot \mathbf{c}_p}{\mathbf{G}_i \cdot \mathbf{c}_p}, \tag{S13}$$

where $\mathbf{G}_i \in \mathbb{R}^{1\times(n_b+n_g)}$ is the $i^{th}$ row vector of $\mathbf{G}$ and analogously for $\mathbf{G}_i'$. The prey specific-uptake rate matrix is given by,

$$\Phi_{g_{i,j}} = \nu^* \varepsilon_{g_i}^2 \left(\frac{c_{p_j}}{c_{p_j} + \frac{1}{\alpha_{\mathbb{S}}}\kappa^* \varepsilon_{g_i}^4}\right)\left(\frac{c_{O_2}}{c_{O_2} + a_{p_j}^C \kappa^* \varepsilon_{g_i}^4}\right) F_T\left(\Delta_r G\left(R_{g_{i,j}}\right), n_{g_{i,j}}^e\right), \tag{S14}$$

where $n_{g_{i,j}}^e = 4a_{p_j}^C$.

### State Space Model

The food web structure, or connectivity, of a system composed of $n_s$ substrates, $n_b$bacteria and $n_g$ consumers is described by the two matrices, $\mathbf{B} \in \mathbb{R}^{n_b \times n_s}$ and $\mathbf{G} \in \mathbb{R}^{n_g \times (n_b + n_g)}$**,** that define which substrates are consumed by each bacteria and which prey each consumer eats, respectively (Fig. 1 in text). To model the experimental chemostats, we use the following state variables: $c_{s_i}$, concentration of carbon substrates in the medium; $c_{b_i}$ concentrations of bacteria; $c_{g_i}$, concentrations of consumers. To model observed experimental variables, we also include: $c_{O_2}$, concentration of dissolved oxygen; $p_{O_2}$, partial pressure of oxygen in bioreactor headspace; $c_{H_2CO_3}$, concentration of all carbonate species; $p_{CO_2}$, partial pressure of $CO_2$ in headspace; $c_{H_3PO_4}$, concentration of inorganic phosphate. All concentrations are in mmol $m^{-3}$, partial pressures are in mbar and nitrogen species are not included as state variables because the experimental system is designed to be P limited. However, ammonium concentration is fixed as a parameter in the model's input file for thermodynamic calculations (a value of 10 µM was used). The differential equations governing the state variables, given the reaction rates defined above, are given by,

*Food web variables*

$$\frac{dc_{s_i}}{dt} = \frac{F_L}{V_L}\left(c_{s_i}^f - c_{s_i}\right) - \sum_{j=1}^{n_b} \frac{r_{b_{j,i}}}{\alpha_{s_i}} \tag{S15}$$

$$\frac{dc_{b_i}}{dt} = \frac{F_L}{V_L}\left(c_{b_i}^f - c_{b_i}\right) + \sum_{j=1}^{n_s} \left(1 - a_{s_j}^A\right) \frac{\varepsilon_{b_i}}{\alpha_{\mathbb{S}}} r_{b_{i,j}} - \sum_{j=1}^{n_g} \frac{r_{g_{j,i}}}{\alpha_{\mathbb{S}}} \tag{S16}$$

$$\frac{dc_{g_i}}{dt} = \frac{F_L}{V_L}\left(c_{g_i}^f - c_{g_i}\right) + \sum_{j=1}^{n_p} \frac{\varepsilon_{g_i} r_{g_{i,j}}}{\alpha_{\mathbb{S}}} - \sum_{j=1}^{n_g} \frac{r_{g_{j,i+n_b}}}{\alpha_{\mathbb{S}}} \tag{S17}$$

*Gases, inorganic carbon, and phosphate*

$$\begin{aligned} \frac{dc_{O_2}}{dt} = \frac{F_L}{V_L}\left(c_{O_2}^f - c_{O_2}\right) + \frac{k_L^{O_2} A}{V_L}\left(p_{O_2} h_{O_2}(T) - c_{O_2}\right) - \sum_{i=1}^{n_b} \sum_{j=1}^{n_s} \left(1 - \varepsilon_{b_i}\right) a_{s_j}^C r_{b_{i,j}} \\ - \sum_{i=1}^{n_g} \sum_{j=1}^{n_p} \left(1 - \varepsilon_{g_i}\right) a_{p_j}^C r_{g_{i,j}} \end{aligned} \tag{S18}$$

$$\frac{dp_{O_2}}{dt} = \frac{F_G}{V_G}\left(p_{O_2}^f - p_{O_2}\right) + \frac{k_L^{O_2} A}{V_G} RT \left(c_{O_2} - p_{O_2} h_{O_2}(T)\right) \tag{S19}$$

$$\begin{aligned} \frac{dc_{H_2CO_3}}{dt} = \frac{F_L}{V_L}\left(c_{H_2CO_3}^f - c_{H_2CO_3}\right) + \frac{k_L^{CO_2} A}{V_L}\left(p_{CO_2} h_{CO_2}(T) - c_{CO_2}\right) \\ + \sum_{i=1}^{n_b} \sum_{j=1}^{n_s} \left(\varepsilon_{b_i} a_{s_j}^A + 1 - \varepsilon_{b_i}\right) r_{b_{i,j}} + \sum_{i=1}^{n_g} \sum_{j=1}^{n_p} \left(1 - \varepsilon_{g_i}\right) r_{g_{i,j}} \end{aligned} \tag{S20}$$

$$\frac{dp_{CO_2}}{dt} = \frac{F_G}{V_G}\left(p_{CO_2}^f - p_{CO_2}\right) + \frac{k_L^{CO_2} A}{V_G} RT \left(c_{CO_2} - p_{CO_2} h_{CO_2}(T)\right) \tag{S21}$$

$$\frac{dc_{H_3PO_4}}{dt} = \frac{F_L}{V_L}\left(c^f_{H_3PO_4} - c_{H_3PO_4}\right) - \sum_{i=1}^{n_b}\sum_{j=1}^{n_s}\left(1 - a^A_{s_j}\right)\frac{\varepsilon_{b_i}\zeta_{\mathbb{S}}}{\alpha_{\mathbb{S}}} r_{b_{i,j}} + \sum_{i=1}^{n_g}\sum_{j=1}^{n_p}\left(1 - \varepsilon_{g_i}\right)\frac{\zeta_{\mathbb{S}}}{\alpha_{\mathbb{S}}} r_{g_{i,j}} \tag{S22}$$

where $F_L$ and $F_G$ are volumetric flow rates (m$^3$ d$^{-1}$) of liquid and gas streams into the bioreactor that have species concentrations of $c_i^f$ and $p_i^f$, respectively, $k_L$ is the liquid-side mass transfer coefficient (aka piston velocity, m d$^{-1}$), $A$ is the total gas-water interface area including bubbles (m$^2$) determined experimentally [4], $V_L$ and $V_G$ are the liquid and gas volumes (m$^3$) in the bioreactor, and $h_i(T)$ is the Henry's law constant (mmol m$^{-3}$ mbar$^{-1}$) for gas $i$ at temperature $T$ (°C) [5]. The set of ODE's are integrated using a Blended implicit Method (BiM) [6].

### Entropy Production

The instantaneous entropy production from irreversible processes, $\dot{\sigma}$ (J K$^{-1}$ d$^{-1}$), occurs as a result of dissipating chemical potential energy stored in substrates, bacteria, and consumers, which is calculated by the product of the reaction free energies, the reaction rate, temperature and system volume. Entropy production from mixing is negligible [7] and is ignored here. The total entropy production associated with bacterial growth and respiration is given by,

$$\dot{\sigma}_{b_T} = -\frac{V_L}{T}\sum_{i=1}^{n_b}\sum_{j=1}^{n_s} r_{b_{i,j}}\Delta_r G\left(R_{b_{i,j}}\right), \tag{S23}$$

and the total entropy production from consumer growth and respiration is similarly given by,

$$\dot{\sigma}_{g_T} = -\frac{V_L}{T}\sum_{i=1}^{n_g}\sum_{j=1}^{n_b+n_g} r_{g_{i,j}}\Delta_r G\left(R_{g_{i,j}}\right), \tag{S24}$$

where $T$ and $V_L$ are temperature and the liquid volume of the bioreactor.

As described in the main text, abiotic systems differ from biological ones in that the latter maximizes entropy production over time, which can be calculated by integrating $\dot{\sigma}_{b_T}$ and $\dot{\sigma}_{g_T}$ over an interval of time, as given by,

$$\langle\dot{\sigma}\rangle\big|_{t_{InvS}}^{\delta_{MEP}} = \frac{1}{\delta_{MEP}} \int_{t_{InvS}}^{t_{InvS}+\delta_{MEP}} \dot{\sigma}_{b_T} + \dot{\sigma}_{g_T} d\tau \tag{S25}$$

Where $\langle\dot{\sigma}\rangle\big|_{t_{InvS}}^{\delta_{MEP}}$ is the average entropy production over $\delta_{MEP}$ days starting at time $t_{InvS}$, where $t_{InvS}$ was set to zero for the simulations in this study.

### Entropy production maximization

Food web dynamics, and associated entropy production, depend on three classes of "traits" or control variables, namely, the growth efficiencies of the bacteria and consumers, $\boldsymbol{\varepsilon}_b$ and $\boldsymbol{\varepsilon}_g$, the elements of the bacterial substate preference matrix, $\mathbf{B}$, and the elements of the consumer prey preference matrix, $\mathbf{G}$. Given the dimensions of these vectors and matrices, the total number of unknowns is $n_u = n_b + n_g + n_b(n_s + n_g) + n_g^2$. To determine the values of the $n_u$ traits, an optimization problem was defined that maximizes entropy production over a specified interval of time while solving the governing set of ODEs, Eqs. (S15 - S22), as given by,

$$J = \max_{\boldsymbol{\varepsilon}_b, \boldsymbol{\varepsilon}_g, \mathbf{B}, \mathbf{G}} \langle\dot{\sigma}\rangle\big|_{t_{InvS}}^{\delta_{MEP}} \tag{S26}$$

This optimization problem was solved for different values of $\delta_{MEP}$ to examine how the timescale changes the diversity and types of substrates bacteria consume (i.e., generalist versus specialists) and the trophic structure of the consumers.

We used a parallel version of BOBYQA [8], hyperBOB [9], that uses a derivative-free optimization approach to find local optima. Because there is no guarantee of finding the global optimum, we ran hyperBOB on a cluster with 200 CPU cores and selected the best solution from the 200 returned.

### Food web metrics

Hill Numbers [10] were used to calculated the substrate diversity metric for bacteria based on the row vectors of the $\mathbf{B}$ matrix as follows,

$$ {}^{q}D_i = \left( \sum_{j=1}^{n_S} \mathbf{B}_{i,j}^{q} \right)^{1/(1-q)} , \quad \text{(S27)} $$

where $q$ was set to 2, so that ${}^{2}D_i$ is an arithmetic mean weighted substrate diversity index. The trophic position of a consumer in the food web, $T_{g_i}$, was calculated from the $\mathbf{G}$ matrix, which is a sub matrix of the full transfer matrix (Fig. S1), as described by Levin [11], which is given by,

$$ T_{g_i} = \sum_{j=1}^{n_b+n_g} \left( \mathbf{I} - \begin{bmatrix} \mathbf{0} \\ \mathbf{G} \end{bmatrix} \right)^{-1}_{n_B+i,j} \quad \text{(S28)} $$

where the zero matrix, $\mathbf{0}$, has the dimensions of $n_b \times (n_b + n_g)$, and the identity matrix, $\mathbf{I}$, is $(n_b + n_g) \times (n_b + n_g)$.

All source code (Fortran) for the model runs is available as a Github repo [12].

### Supplemental Figures

**Source** (columns) / **Consumes** (rows)

| Consumes \ Source | Glucose | Acetate | Ethanol | Methanol | Xylose | b1 | b2 | b3 | b4 | b5 | b6 | b7 | b8 | b9 | b10 | g1 | g2 | g3 | g4 | g5 |
|---|---|---|---|---|---|---|---|---|---|---|---|---|---|---|---|---|---|---|---|---|
| **Glucose** | 1 | 0 | 0 | 0 | 0 | 0 | 0 | 0 | 0 | 0 | 0 | 0 | 0 | 0 | 0 | 0 | 0 | 0 | 0 | 0 |
| **Acetate** | 0 | 1 | 0 | 0 | 0 | 0 | 0 | 0 | 0 | 0 | 0 | 0 | 0 | 0 | 0 | 0 | 0 | 0 | 0 | 0 |
| **Ethanol** | 0 | 0 | 1 | 0 | 0 | 0 | 0 | 0 | 0 | 0 | 0 | 0 | 0 | 0 | 0 | 0 | 0 | 0 | 0 | 0 |
| **Methanol** | 0 | 0 | 0 | 1 | 0 | 0 | 0 | 0 | 0 | 0 | 0 | 0 | 0 | 0 | 0 | 0 | 0 | 0 | 0 | 0 |
| **Xylose** | 0 | 0 | 0 | 0 | 1 | 0 | 0 | 0 | 0 | 0 | 0 | 0 | 0 | 0 | 0 | 0 | 0 | 0 | 0 | 0 |
| **b1** | 0.36 | 0.00 | 0.60 | 0.00 | 0.04 | 0 | 0 | 0 | 0 | 0 | 0 | 0 | 0 | 0 | 0 | 0 | 0 | 0 | 0 | 0 |
| **b2** | 0.42 | 0.11 | 0.28 | 0.04 | 0.16 | 0 | 0 | 0 | 0 | 0 | 0 | 0 | 0 | 0 | 0 | 0 | 0 | 0 | 0 | 0 |
| **b3** | 0.29 | 0.00 | 0.59 | 0.00 | 0.12 | 0 | 0 | 0 | 0 | 0 | 0 | 0 | 0 | 0 | 0 | 0 | 0 | 0 | 0 | 0 |
| **b4** | 0.10 | 0.00 | 0.62 | 0.00 | 0.29 | 0 | 0 | 0 | 0 | 0 | 0 | 0 | 0 | 0 | 0 | 0 | 0 | 0 | 0 | 0 |
| **b5** | 0.00 | 0.14 | 0.32 | 0.54 | 0.00 | 0 | 0 | 0 | 0 | 0 | 0 | 0 | 0 | 0 | 0 | 0 | 0 | 0 | 0 | 0 |
| **b6** | 0.00 | 0.15 | 0.00 | 0.85 | 0.00 | 0 | 0 | 0 | 0 | 0 | 0 | 0 | 0 | 0 | 0 | 0 | 0 | 0 | 0 | 0 |
| **b7** | 0.36 | 0.20 | 0.00 | 0.00 | 0.43 | 0 | 0 | 0 | 0 | 0 | 0 | 0 | 0 | 0 | 0 | 0 | 0 | 0 | 0 | 0 |
| **b8** | 0.21 | 0.00 | 0.53 | 0.00 | 0.26 | 0 | 0 | 0 | 0 | 0 | 0 | 0 | 0 | 0 | 0 | 0 | 0 | 0 | 0 | 0 |
| **b9** | 0.20 | 0.02 | 0.24 | 0.34 | 0.20 | 0 | 0 | 0 | 0 | 0 | 0 | 0 | 0 | 0 | 0 | 0 | 0 | 0 | 0 | 0 |
| **b10** | 0.13 | 0.00 | 0.61 | 0.00 | 0.26 | 0 | 0 | 0 | 0 | 0 | 0 | 0 | 0 | 0 | 0 | 0 | 0 | 0 | 0 | 0 |
| **g1** | 0 | 0 | 0 | 0 | 0 | 0.30 | 0.01 | 0.00 | 0.00 | 0.00 | 0.00 | 0.01 | 0.28 | 0.01 | 0.00 | 0.01 | 0.06 | 0.32 | 0.00 | 0.00 |
| **g2** | 0 | 0 | 0 | 0 | 0 | 0.10 | 0.00 | 0.03 | 0.00 | 0.23 | 0.22 | 0.00 | [illegible] | 0.00 | 0.02 | 0.00 | 0.00 | 0.22 | 0.06 | 0.06 |
| **g3** | 0 | 0 | 0 | 0 | 0 | 0.12 | 0.09 | 0.01 | 0.12 | 0.05 | 0.12 | 0.01 | 0.06 | 0.12 | 0.03 | 0.09 | 0.05 | 0.00 | 0.01 | 0.11 |
| **g4** | 0 | 0 | 0 | 0 | 0 | 0.00 | 0.00 | 0.03 | 0.00 | 0.04 | 0.07 | 0.00 | [illegible] | 0.00 | 0.02 | 0.00 | 0.00 | 0.84 | 0.00 | 0.00 |
| **g5** | 0 | 0 | 0 | 0 | 0 | 0.00 | 0.00 | 0.00 | 0.23 | 0.28 | 0.28 | 0.00 | 0.00 | 0.00 | 0.09 | 0.00 | 0.00 | 0.11 | 0.00 | 0.00 |

Block labels: **I** (substrates × substrates); **0** Cross feeding (substrates × b1–g5); **B** (bacteria × substrates); **0** Bacterial Predators Pathogens (bacteria × b1–g5); **0** Mixotroph (g1–g5 × substrates); **G** (g1–g5 × b1–g5).

**Fig. S1**. Full transfer matrix for a system of 5 substrates, 10 bacteria, and 5 consumers. In the model implemented for this study, it was assumed that there were no cross-feeding, bacterial predators or pathogens, nor mixotrophs. The diagonal purple line on the right side of the $\mathbf{G}$ matrix highlights cannibalism. The identity matrix, $\mathbf{I}$, in the upper left of the transfer matrix signifies that the substrate added went to that substrate. In Levine's [11] transition matrix, his sub matrix $\mathbf{R}$ equals $\begin{bmatrix}\mathbf{B}\\ \mathbf{0}\end{bmatrix}$ and $\mathbf{Q}$ equals $\begin{bmatrix}\mathbf{0}\\ \mathbf{G}\end{bmatrix}$.

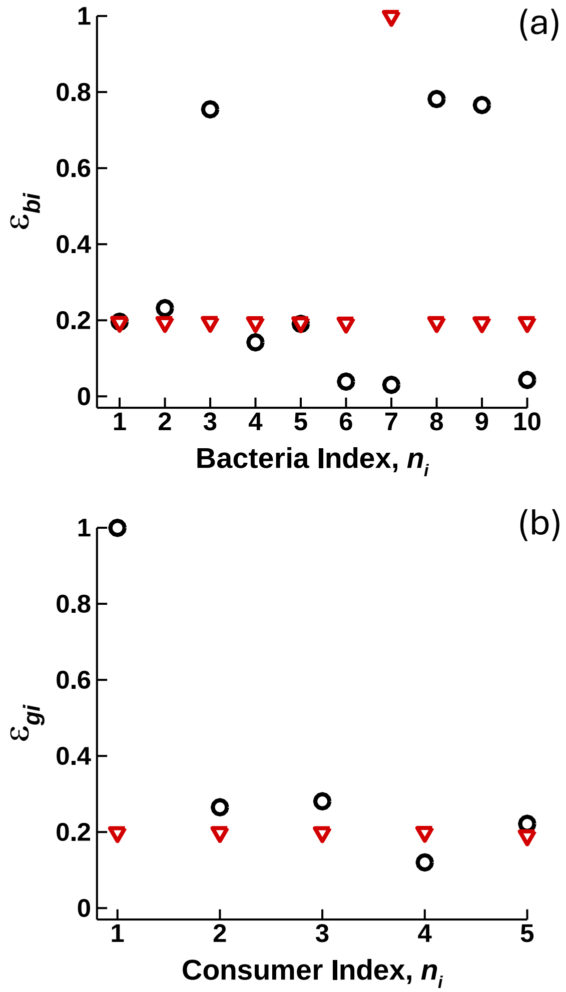


**Fig. S2**. Growth efficiencies of (a) bacteria, $\varepsilon_{b_i}$, and (b) consumers, $\varepsilon_{b_i}$, for the top ranked short-term optimization (STO) solution (red triangles) and the long-term optimization (LTO) solution (black circles)

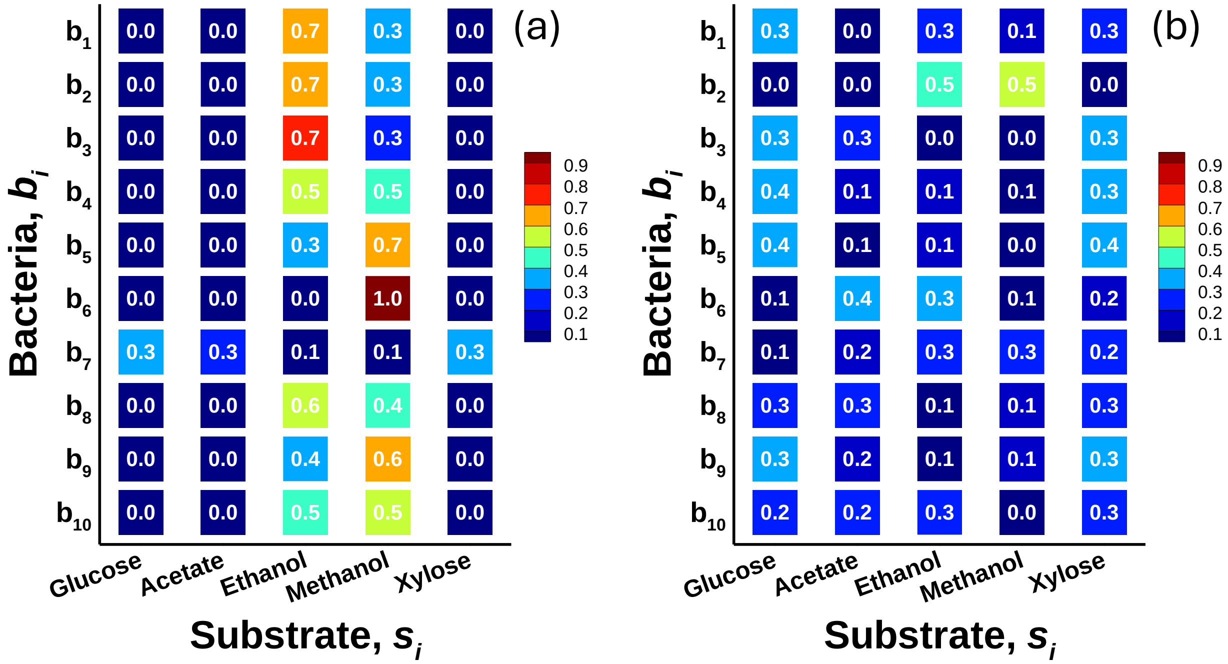


**Fig. S3**. Substrate preference matrix of bacteria, $\mathbf{B}$, for the top-ranked (a) short- term optimization (STO) solution and (b) long-term optimization (LTO) solution.

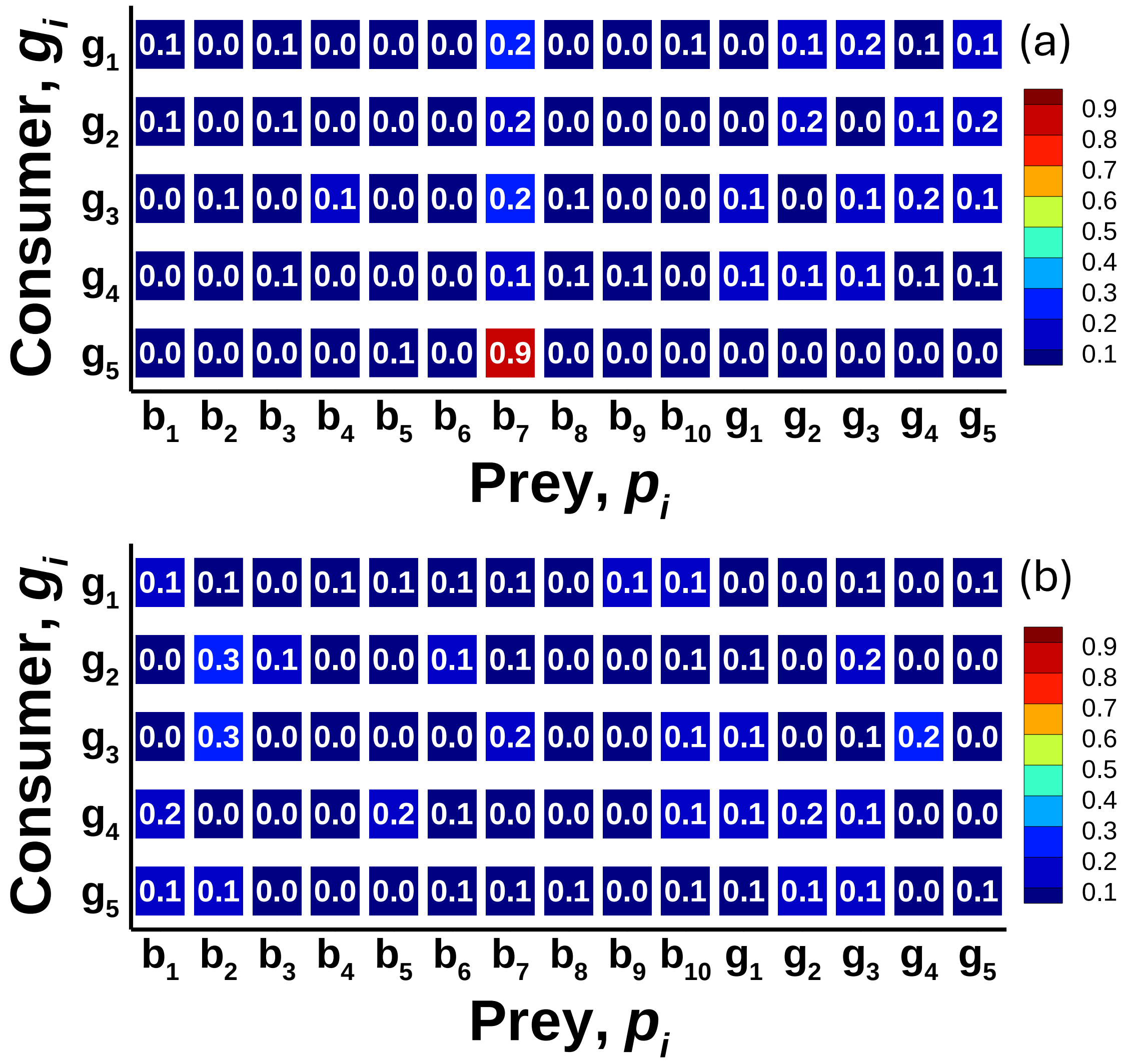


**Fig. S4**. Prey preference matrix of consumers, $\mathbf{G}$, for the top-ranked (a) short-term optimization (STO) solution and (b) long-term optimization (LTO) solution.

### Input files for the Short-Term Optimization (STO) and Long-Term Optimization (LTO) simulations

**Run43, STO simulation:**

```
! 11-Apr-2025
!   This is Run20, but using V2.2d instead of V2.2c
!   Darwin_ChemoStat_V2.2d.f90
!      Now effective trophic position of the consumers is calculated.
!   2 day optimization window
!   1 uM P
!   D = 0.1 1/d (F_L = 0.0003 )

&dimens
! problem dimensions
nSub = 5
nBac = 10
nGrz = 5
/

&parameters
! All parameters used by problem
absZero = 1.d-8  ! prevents division by zero

kappa = 5000. ! (mmole/m3)
nuStar = 350. ! (1/d)
delPsi = 0.1 !0.240 ! LaRowe's thermo driver.  Cell membrane potential (V)

alf_Bio = 1.000 ! C elemental composition of phytoplankton, unit carbon based. (use
Battley1998 for yeast)
bet_Bio = 1.613 ! H
gam_Bio = 0.557 ! O
del_Bio = 0.158 ! N
zet_Bio = 0.012 ! P

T_K = 293.15 ! temperature (K)
pH = 6.6     ! pH
is = 0.07    ! ionic strength (M)  Can use the formula: 0.72*PSU/35.0 (assume 3.4 PSU
here)

nh3_hold = 10.0 ! Concentration (mmol/m3) of NH3 used in thermo calculations

kL_o2  = 1.5 ! piston velocity for O2 (m/d)
kL_co2 = 1.3 ! piston velocity for CO2 (m/d)  Should use Schmidt no. to relate these,
do it later
area_GL = 0.041 ! area of gas-liquid interface (m2) (MC's are about 9" in diameter)
V_L = 0.003 ! volume of liquid in chemostat (m3)
F_L = 0.0003 ! liquid flow rate (m3/d)
V_G = 0.00474 ! volume of gas headspace (m3)
F_G = 0.01545 ! gas flow rate (m3/d) (This is 10 sccm at 273 K, or 10.73 mL/min at 20
C)

! substrate names
subNames(1) = 'glucose'
subNames(2) = 'acetate'
subNames(3) = 'ethanol'
subNames(4) = 'methanol'
subNames(5) = 'xylose'

! substrate CHO composition
CHO(1,:) = 6, 12, 6
CHO(2,:) = 2,  4, 2
CHO(3,:) = 2,  6, 1
```

```
CHO(4,:) = 1,  4, 1
CHO(5,:) = 5, 10, 5

! initial conditions for all bi and gi (mmol/m3)
! The chemostat feed is set to these initial conditions, except no bi or gi in it.
t0 = 0.             ! start time. (d)
tDays   = 53. ! number of days to run
t0_mep = 0. ! the time (day) that entropy production maximization begins
tDays_mep = 2. ! the number of days over which entropy production is maximized
bi_ini = 0.1
bi_f   = 0.0  ! No bacteria in feed
gi_ini = 0.1
gi_f   = 0.0  ! No grazers in feed.
! substrates (mmol/m3)
sj_f(1) = 50.0 ! glucose
sj_f(2) = 167.5 ! acetate
sj_f(3) = 108.3 ! ethanol
sj_f(4) = 206.1 ! methanol
sj_f(5) = 59.69 ! xylose
! Chemistry (mmol/m3 and mbar)
chm_f(1) = 250      ! oxygen
chm_f(2) = 212.7    ! O2 pressure (mbar)
chm_f(3) = 210      ! DIC (this one needs some thought, but currently 0.1 of seawater)
chm_f(4) = 0.405    ! pCO2 (mbar)
chm_f(5) = 1.       ! phosphate, if using phosphate buffer.
/

&hyperBOBparams
! hyperBOB parameters
optimize = .true. ! if set to false then traits values are randomly set and a
simulation is run. Mostly for testing
rhobeg = 0.49 ! initial and final values of a trust region radius
rhoend = 0.0001
iprint = 0  ! controls amount of printing (0, 1, 2 or 3)
maxfun = 200000  ! maximum number of calls to CALFUN
fcnUpdate = 100 ! After every fcnUpdate PDE integration, info is printed out.
seed_bobyqa = 17 ! used by hyperBOB for random hypercube. All MPI processes use same
value
/

&tecIOparams
! TecIO parameters for output
nIOpts = 1000 ! number of time series points to store in a single zone
debug_IO = 0 ! 1 to debug, 0 no debug.
flushData = .false. ! If true, data is flushed to files at each time output, but may
slow down execution.
/

&BiMparams
! parameters for BiM integrator
rtol = 1.d-6 ! relative tolerance
atol = 1.d-8 ! absolute tolerance
hmax_BiM = 0.1 ! maximum step size.  = (tend-t0)/8 if set to 0
maxstep_BiM = 0 ! maximum number of steps. Default 100000 if set to zero
maxattempts = 10 ! maximum number of times to retry integration
ompThreads = 1 ! Number of OMP threads to use for numerical Jacobian calculations
minCompFac = 100000.0 ! if a ODE solution takes longer than (tf-t0)/minCompFac (days),
then it is terminated.
/
```

**Run44, LTO simulation:**

Same as above, except:

```
tDays_mep = 53. ! the number of days over which entropy production is maximized
```
